# Metabolic Alteration in Oxylipins and Endocannabinoids Point to an Important Role for Soluble Epoxide Hydrolase and Inflammation in Alzheimer’s Disease - Finding from Alzheimer’s Disease Neuroimaging Initiative

**DOI:** 10.1101/2025.04.01.646677

**Authors:** Kamil Borkowski, Chunyuan Yin, Alida Kindt, Nuanyi Liang, Elizabeth de Lange, Colette Blach, John W. Newman, Rima Kaddurah-Daouk, Thomas Hankemeier, Alzheimer’s Disease Metabolomics Consortium, the Alzheimer’s Disease Neuroimaging Initiative

**Affiliations:** West Coast Metabolomics Center, Genome Center, University of California Davis, Davis, CA 95616, USA; Metabolomics and Analytics Centre, Leiden Academic Center for Drug Research, Leiden University, The Netherlands; Research Division of Systems Pharmacology and Pharmacy, LACDR, Leiden University, The Netherlands; Department of Psychiatry and Behavioral Sciences, Duke Institute for Brain Sciences and Department of Medicine, Duke University, Durham, NC 27708, USA; Western Human Nutrition Research Center, United States Department of Agriculture - Agricultural Research Service, Davis, CA 95616, USA; Department of Nutrition, University of California - Davis, Davis, CA 95616, USA; Predictive Pharmacology, Division of Systems Pharmacology and Pharmacy, Leiden Academic Center for Drug Research, Leiden University, The Netherlands

**Keywords:** Metabolomics, lipidomics, inflammation, mild cognitive impairment, Alzheimer’s Disease, oxylipins, endocannabinoids, soluble epoxide hydrolase

## Abstract

Mounting evidence implicates inflammation as a key factor in Alzheimer’s disease (AD) development. We previously identified pro-inflammatory soluble epoxide hydrolase (sEH) metabolites to be elevated in plasma and CSF of AD patients and to be associated with lower cognition in non-AD subjects. Soluble epoxide hydrolase is a key enzyme converting anti-inflammatory epoxy fatty acids to pro-inflammatory diols, reported to be elevated in multiple cardiometabolic disorders. Here we analyzed over 700 fasting plasma samples from the baseline of Alzheimer’s Disease Neuroimaging Initiative (ADNI) 2/GO study. We applied targeted mass spectrometry method to provide absolute quantifications of over 150 metabolites from oxylipin and endocannabinoids pathway, interrogating the role for inflammation/immune dysregulation and the key enzyme soluble epoxide hydrolase in AD. We provide further insights into the regulation of this pathway in different disease stages, APOE genotypes and between sexes. Additionally, we investigated in mild cognitive impaired (MCI) patients, metabolic signatures that inform about resilience to progression and conversion to AD. Key findings include I) confirmed disruption in this key central pathway of inflammation and pointed to dysregulation of sEH in AD with sex and disease stage differences; II) identified markers of disease progression and cognitive resilience using sex and ApoE genotype stratified analysis highlighting an important role for bile acids, lipid peroxidation and stress response hormone cortisol. In conclusion, we provide molecular insights into a central pathway of inflammation and links to cognitive dysfunction, suggesting novel therapeutic approaches that are based on targeting inflammation tailored for subgroups of individuals based on their sex, APOE genotype and their metabolic profile.

## Introduction

Inflammation is recognized as an important contributor to Alzheimer’s disease (AD) pathology and cognitive decline and as a crucial risk factor for AD development that exacerbates disease progression (1, 2) .Oxylipin and endocannabinoid (OxyL-EC) pathways are key regulators of inflammation, with changes in their levels being a hallmark of inflammation-related disorders (3–13). The oxylipins including fatty acid alcohols, diols, epoxides, ketones, and prostanoids are derived from multiple polyunsaturated fatty acids (PUFA) by the action of cyclooxygenases (COX), lipoxygenases (LOX), cytochrome P450 (CYP), soluble epoxide hydrolase (sEH) or reactive oxygen species (ROS) and various downstream enzymatic processes (14). Circulating endocannabinoids are produced either by acylation and release of acyl ethanolamides from phosphatidylethanolamine and further degraded by fatty acid amide hydrolase (FAAH), or as a product of glycerol-lipid metabolism like monoacylglycerols (MAGL) (15).

Increasing evidence has demonstrated OxyL-EC dysregulation in AD patients. Disturbances at the protein expression and metabolome level of these pathways are observed in both the periphery (16, 17) and central nervous system (18–20) in humans. In rodent models of AD, AD-related pathologies can be altered by manipulating OxyL-EC related enzymes, including 5-LOX (21, 22), 12/15-LOX (23–25), COX (26, 27), sEH (28, 29), CYP (30, 31), FAAH (32), MAGL (33), and their combination (34). Moreover, genetic polymorphisms in many of these pathways have been linked to AD risk, supporting causative effects from these pathways (35–37).

The Alzheimer’s Disease Metabolomics Consortium (ADMC) has shown that peripheral metabolic changes informed about cognitive changes, brain imaging changes, and ATN markers for disease (38–41). We have recently generated a comprehensive profile of OxyL-EC pathways in subsets of well-characterized AD cohorts, including the Religious Orders Study and the Rush Memory and Aging Project (ROS-MAP) (42) and the Emory cohort (the Emory Healthy Brain Study, Cognitive Neurology Research, and Memory) (43). We have used a state-of-the-art validated, quantitative, targeted mass spectrometry platform, providing absolute quantification of ∼ 150 OxyL-EC, covering multiple metabolic pathways and possible substrates (44, 45). This pioneer work identified metabolites of CYP/sEH and acyl ethanolamides in plasma and CSF to be associated with AD pathology, and plasma sEH metabolites to be associated with perceptual speed in cognitively normal and mild cognitively impaired subjects. Furthermore, utilizing the metabolomic-proteomic integration approach, we have shown association of the sEH pathway with CSF proteins related to glycolysis, vascular inflammation and neuronal outgrowth (46), with changes in CSF levels of those proteins being reflective of AD-related changes in the brain (47).

In the current manuscript we seek to confirm and expand on our previous findings of the dysregulation in OxyL-EC pathways in AD, utilizing 763 subjects from the Alzheimer’s Disease Neuroimaging Initiative (ADNI) 2/GO cohort. The ADNI cohort enables us to further investigate the dysregulation of OxyL-EC pathways at different disease stages, ranging from cognitively normal (CN), mild cognitively impaired (MCI) to AD, and the association of baseline OxyL-EC levels with future cognitive decline. Additionally, to uncover disease heterogeneity, we investigate the influence of sex and APOE genotype on the OxyL-EC interaction with AD-related outcomes.

## Methods

### Study Participants

The Alzheimer’s Disease Neuroimaging Initiative (ADNI) study recruited 763 individuals over the age of 70 years into longitudinal study. At baseline (the time when patients enrolled in the project), this comprised 178 cognitively normal controls (CN), 445 with mild cognitive impairment (MCI; 290 early-stage and 155 late-stage MCI) and 136 with Alzheimer’s Disease (AD). The early-stage and late-stage MCI patients were grouped together as the distinction between these two groups is minimal.

Follow-up cognition tests were performed at 1, 2 and 4 years after baseline. We analyzed 763 plasma samples from baseline, from which 23 non-fasting participants were excluded. Additional participant group classification was based on the experienced diagnostic change, i.e., progression from CN to MCI or AD, or from MCI to AD within 1, 2 or 4 years after baseline. “Stable CN” comprises CN baseline samples that did not convert to either MCI or AD within 4 years follow up, and “stable MCI” comprises MCI baseline patients that did not convert to AD within 2 years follow up. Patients whose diagnosis reverted, i.e., changed from AD to MCI or CN, or from MCI to CN were excluded (n=39). See Table-1 and Table-2 for more details.

**Table-1.**
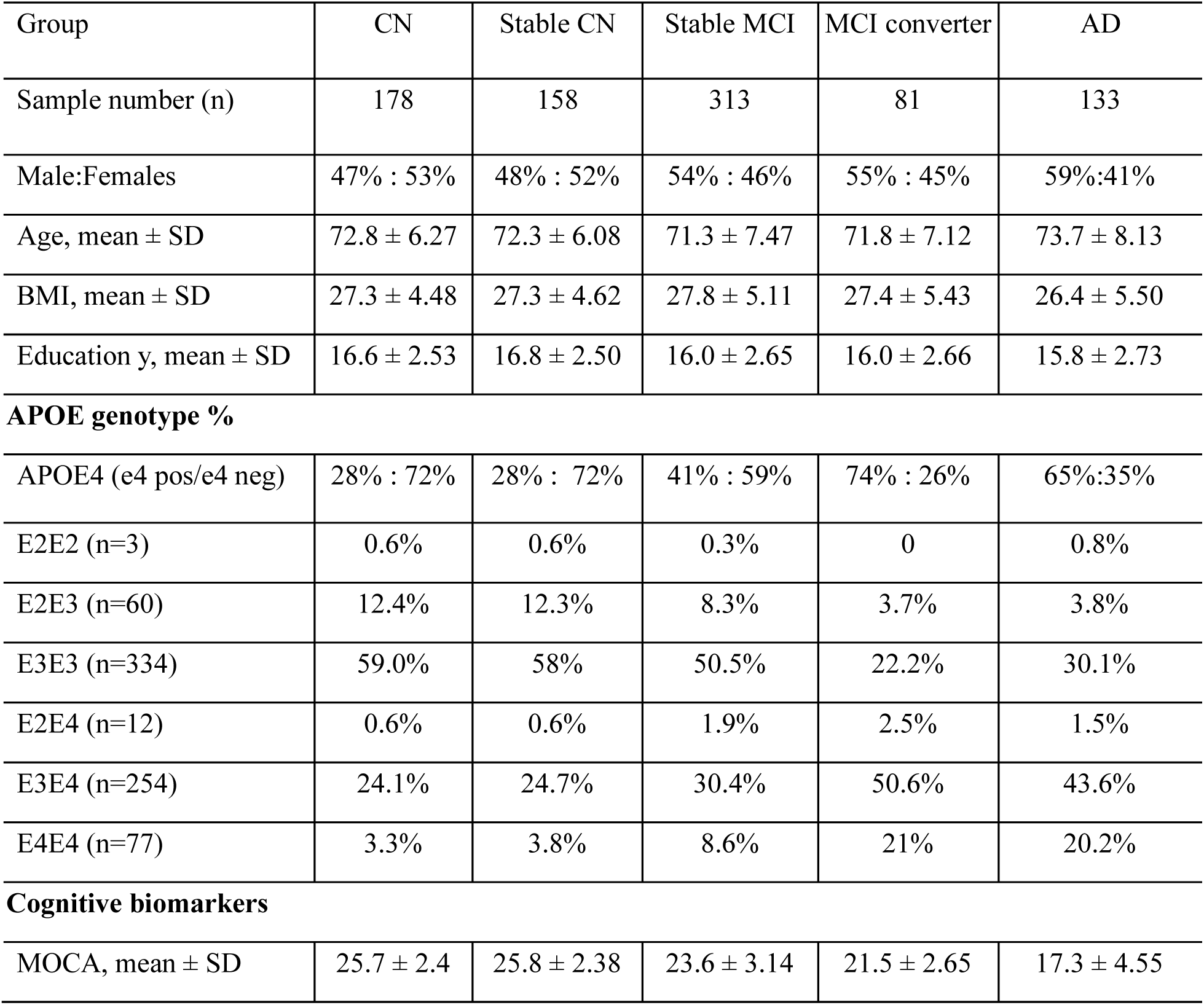
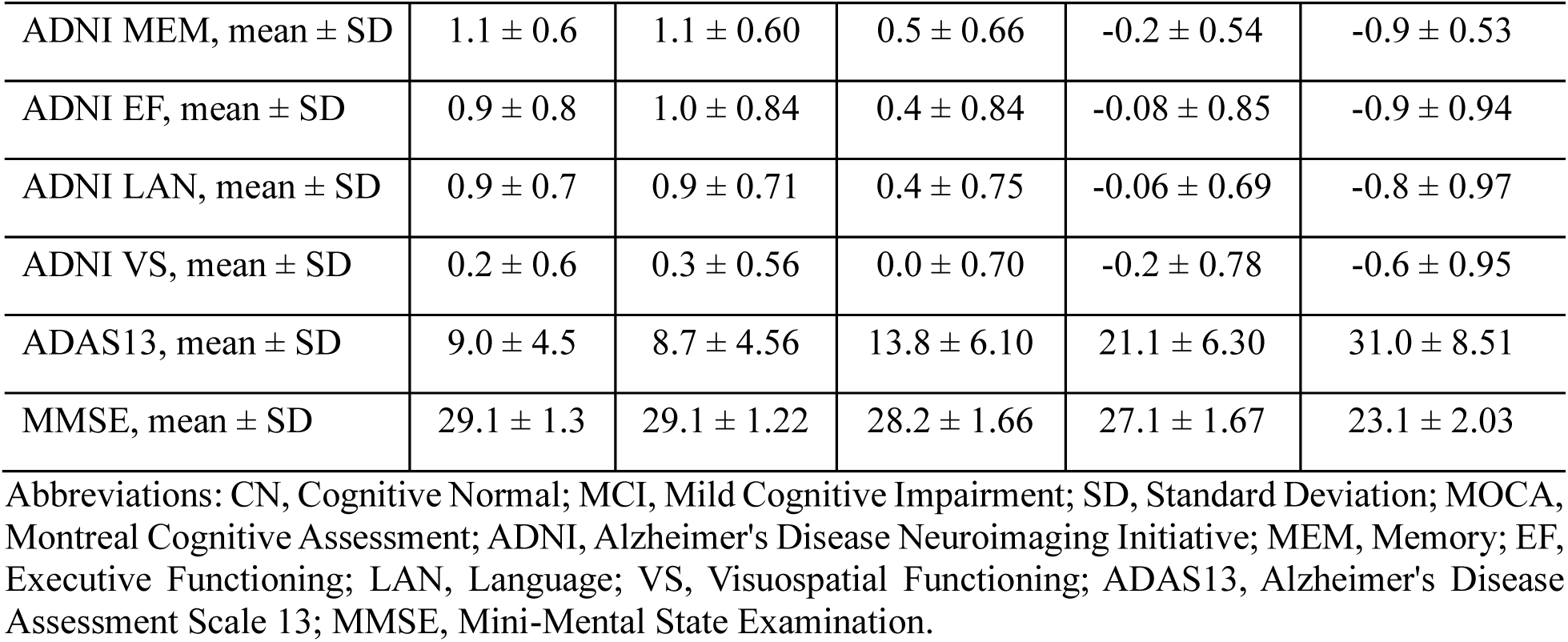
Demographics and clinical data of studied ADNI subjects at baseline.

**Table-2.** Diagnoses for subjects with available CSF measurements in each visit.

|  | **Baseline CN** | **Baseline MCI** | **Baseline AD** |
| --- | --- | --- | --- |
| **Baseline** | 178 CN | 429 MCI | 133 AD |
| **2 years** | 137 CN, 13 MCI, 2 AD, 26 NA | 19 CN, 266 MCI, 60 AD, 84 NA | 1 MCI, 28 AD, 104 NA |
| **4 years** | 94 CN, 11 MCI, 2 AD, 71 NA | 27 CN, 156 MCI, 48 AD, 198 NA | 133 NA |

Abbreviations: AD, Alzheimer's disease; CN, Cognitive Normal; MCI, Mild Cognitive Impairment; NA, cognitive test result not available

### Metabolomics profiling

All baseline plasma samples (n=763) were assessed using a targeted UHPLC-MS/MS lipidomic profiling strategy as previously described (44). Briefly, liquid-liquid extraction with 1:1 methyl tertbutyl ether/n-butanol was performed on plasma samples after including isotopically labeled internal standards and targets were measured using reversed-phase liquid chromatography-mass spectrometry (RPLC-MS/MS) under high and low pH conditions after solvent exchange. Specific UHPLC systems and columns were employed for optimal separation and detection and data preprocessing was performed with vendor-specific software to ensure accurate peak integration and internal standard matching. Briefly, for the high pH measurements a SCIEX 6500+ QTRAP mass spectrometer was used with a Kinetex EVO column by Phenomenex followed by analyst data acquisition software (Version V1.7.2, AB Sciex). The triple quadrupole mass spectrometer operated in polarity switching mode and all analytes were monitored in dynamic Multiple Reaction Monitoring (dMRM) mode. While for the low pH analysis, a Sciex 7500 QTRAP mass spectrometer was used with an Acquity UPLC BEH C18 column (Waters) followed by Sciex OS Software V2.0.0.45330 (AB Sciex). This platform enables analyses of 260 metabolites covering oxylipins and their free fatty acid precursors, lysophospholipids, sphingoid bases, PAF, endocannabinoids and bile acids. All samples were processed with rigorous quality control measures including case/control randomization over all batches, inclusion of solvent blanks, pooled sample quality control aliquots, and NIST Standard Reference Material 1950 – Metabolites in Human Plasma (Merck, Darmstadt, Germany). These batches were re-randomized for measurements where method blanks, reference and solvent calibration samples were included at regular intervals. The majority of analytes were quantified against analytical standards, where area counts were recorded, adjusted for deuterated surrogate responses and then calculated into nanomolar concentrations using solvent calibration lines from which the lowest calibration point was subtracted from other calibration points. Next, the calibration lines were checked for outliers which were identified using an outlier test, where only calibration points at the ends were allowed to be removed. A calibration point within the calibration line was removed if the ratio was lower than its previous (lower) calibration point. After these corrections, the model was recalculated. Reported monoacylglycerols (MAGs) are the sum of 1- and 2-acyl isomers and 10-Nitrooleate is the sum of 9-Nitrooleate and 10-Nitrooleate, due to isomerization during sample processing.

### Data preprocessing

After peak integration an inhouse quality assessment tool was used to perform correction of target compounds with internal standards to obtain relative ratios, batch correction, and filtering of targets with low signal to noise ratios. To obtain absolute concentrations, linear regression models were calculated per batch between known concentrations and relative ratios for each target. The ratio of the lowest calibration point (CAL0) was subtracted from all ratios in the calibration line. Baseline data with detectable coverage of over 70% were further analyzed. All data were log2-transformed. The following data pre-processing procedure was performed in JMP® Pro 17.2.0 (SAS Institute Inc, Cary NC): outliers were detected and removed using the Robust Fit Outliers method and missing values were imputed with the Multivariate Normal Imputation method (48). Ratios of metabolites were calculated based on the base-2 exponential-transformed data after imputation, followed by log2-transformation before further analysis. Non-fasted participants (n =23) and participants of Significant Memory Concern (SMC) were excluded from further analysis. Medication adjustments were made based on the stepwise AIC-backward method previously described (49, 50).

## Statistical analysis

### Data reduction

To reduce the dimensionality and collinearity of the data, gain additional information about associations among variables and to facilitate result interpretation variables showing significant associations with the outcomes (Determined by ANOVA) were converted into cluster components. Variables were clustered using the Jmp Pro implementation of the VARCLUS Procedure, a principal components analysis for variable grouping and cluster component (the linear combination of all variables in each cluster) calculation. This is a data-driven approach where the algorithm defines the optimal number of clusters, with resulting clusters composed of highly correlated variables. Those correlations in terms of metabolomic data mostly align with biological pathways, further confirming integrity of the data.

### Comparison of lipid mediator levels between diagnosis groups

For lipid mediator levels, ANOVA followed by Dunnett’s post hoc test was performed to compare means between CN and other diagnosis groups, stratified by sex. Lipid mediators exhibiting significant differences between groups were grouped into clusters as described above. Cluster components were used in ANOVA analysis with Dunnett post-test, comparing the difference in lipid mediator levels between CN and other diagnosis groups. **Supplemental Table S1 and Supplemental Table S2** describe cluster members as well as their correlations within the cluster, least square means (standard error range) and ANOVA p-values. ANOVA p-values for individual metabolites are presented in the **Supplemental Table S3**. Presence of the APOE 4 allele, sex, body mass index (BMI), age and years of education were included as covariates in all analyses. This analysis was performed in R studio software (Version 4.3.1). Figures were created in R, cytoscape 3.10.2. Multiple comparison control was accomplished with the false discovery rate (FDR) correction method of Benjamini and Hochberg (51)

### MCI conversion and memory (ADNI MEM) predictive minimal models

Using JMP 17.0 pro (JMP, SAS institute, Cary, NC), we applied a combination of bootstrap forest and stepwise linear (for ADNI MEM) and logistic (for MCI converters vs stable MCI) regression modeling, with Bayesian Information Criterion (BIC) cutoff. Variable selection by bootstrap forest minimized outlier effects. Variables most frequently appearing in the models were identified by bootstrap forest (logistic or regression, respectively): trees in forest =100; terms sampled per split =5; bootstrap sample rate =1. A variable contribution scree plot was generated using variable rank and the likelihood ratio of chi-square (for categorical fasted/non-fasted prediction) or sum of squares (for continues cognitive scores). The scree plot was used to determine a likelihood ratio of chi-square or sum of squares cutoff value for variables contributing to the model. Selected variables were then subjected to forward stepwise logistic or linear regression modelling. Stepwise regression was used to highlight independent predictors of outcomes, to create the minimal model.

## Results

In the current study we have analyzed plasma from the 763 participants from the ADNI 2/GO, providing absolute quantification of 45 eicosanoids, 5 endocannabinoids, 49 lysophospholipids, 13 monounsaturated and polyunsaturated fatty acid (MUFA&PUFA), 4 sphingolipids, and 1 steroids lipids (**Figure 1 and Supplemental Table S3**). In addition, our data included 18 bile acids (**Supplemental Figure S1**). The bile acid levels in ADNI cohort were previously reported (40) and are used in the current analysis only to show their interaction with lipid mediators.

**Figure 1.**
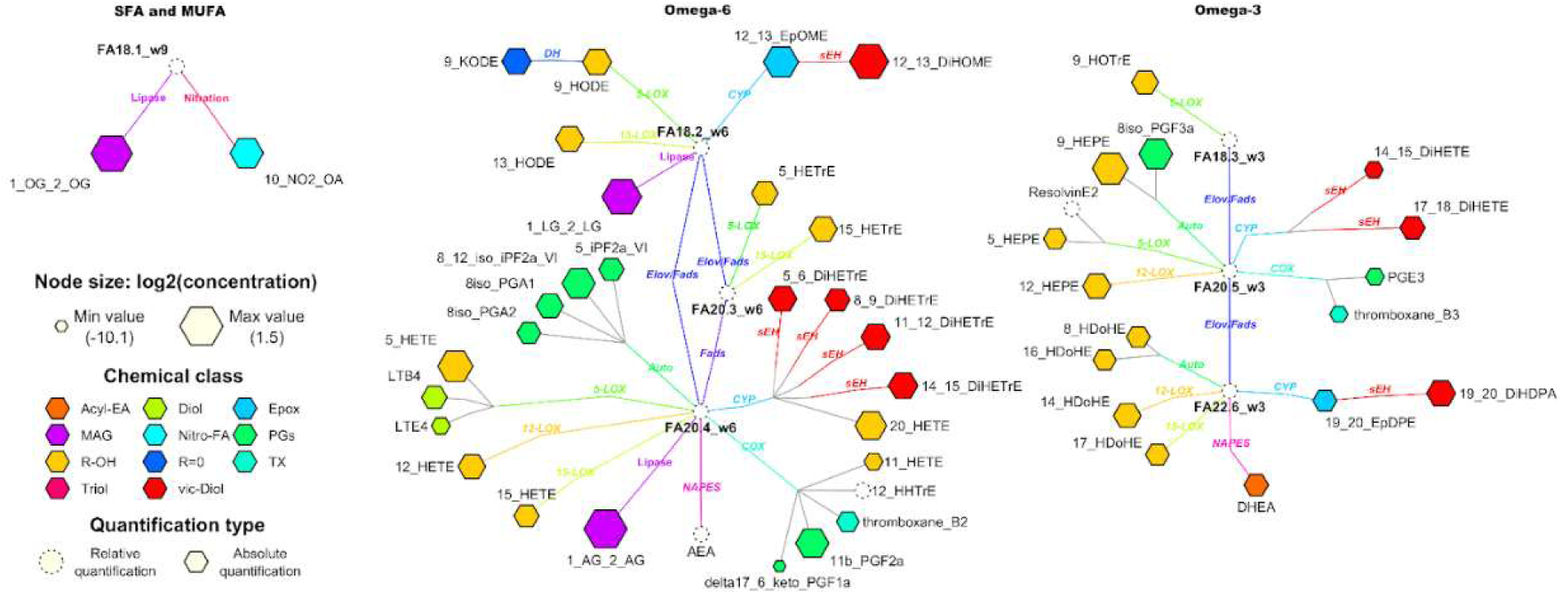
Quantified oxylipins, endocannabinoids and polyunsaturated fatty acids in plasma samples of ADNI 2/GO participants, projected onto their metabolic pathway. Nod size indicates the average concentration observed across the samples, with chemical classes indicated by node color.

### 1. Sex-specific differences in sEH metabolites along disease trajectories

To assess the changes in oxylipin metabolism along disease trajectory, we compared the differences in their levels between diagnosis groups, in the analysis stratified by sex. The following groups were used: cognitively normal (CN), mild cognitively impaired (MCI) and dementia (AD). We excluded subjects who were cognitively normal at baseline who change their status at the 4-year follow-up. The differences between the groups were compared using ANOVA with significant metabolites further converted into cluster components (**Figure 2**). Factorial analysis with sex x group interaction was applied to identify metabolites affected differently by diagnosis group between males and females. Out of 224 metabolites and informative ratios and summations, 21 showed differences between groups. The 21 significant variables were condensed into 6 cluster components by variable clustering and reanalyzed by ANOVA with a Dunnett post-test against CN in a sex-stratified analysis (**Figure 2**). Individual cluster members, their contribution to the cluster and their mean differences between groups are described in the **Supplemental Table S1 and Supplemental Table S2**. The main differences were observed in fatty acid vicinal diols (products of sEH metabolism, Clusters 1 for eicosapentanoic acid (EPA) and docosahexanoic acid (DHA) and Cluster 2 for the arachidonic acid (AA) derivatives) with sex differences. In Females, both omega-3 (EPA and DHA) and omega-6 (AA) derivatives were higher in AD when compared to the CN. However, the MCI was not different from the CN. Additionally, differences were observed in the levels of DHA-derived LOX metabolites (HDoHEs) and LPAs (Clusters 3 and 4) with MCI showing higher levels than CN and cortisol with AD showing higher levels than HC.

**Figure 2.**
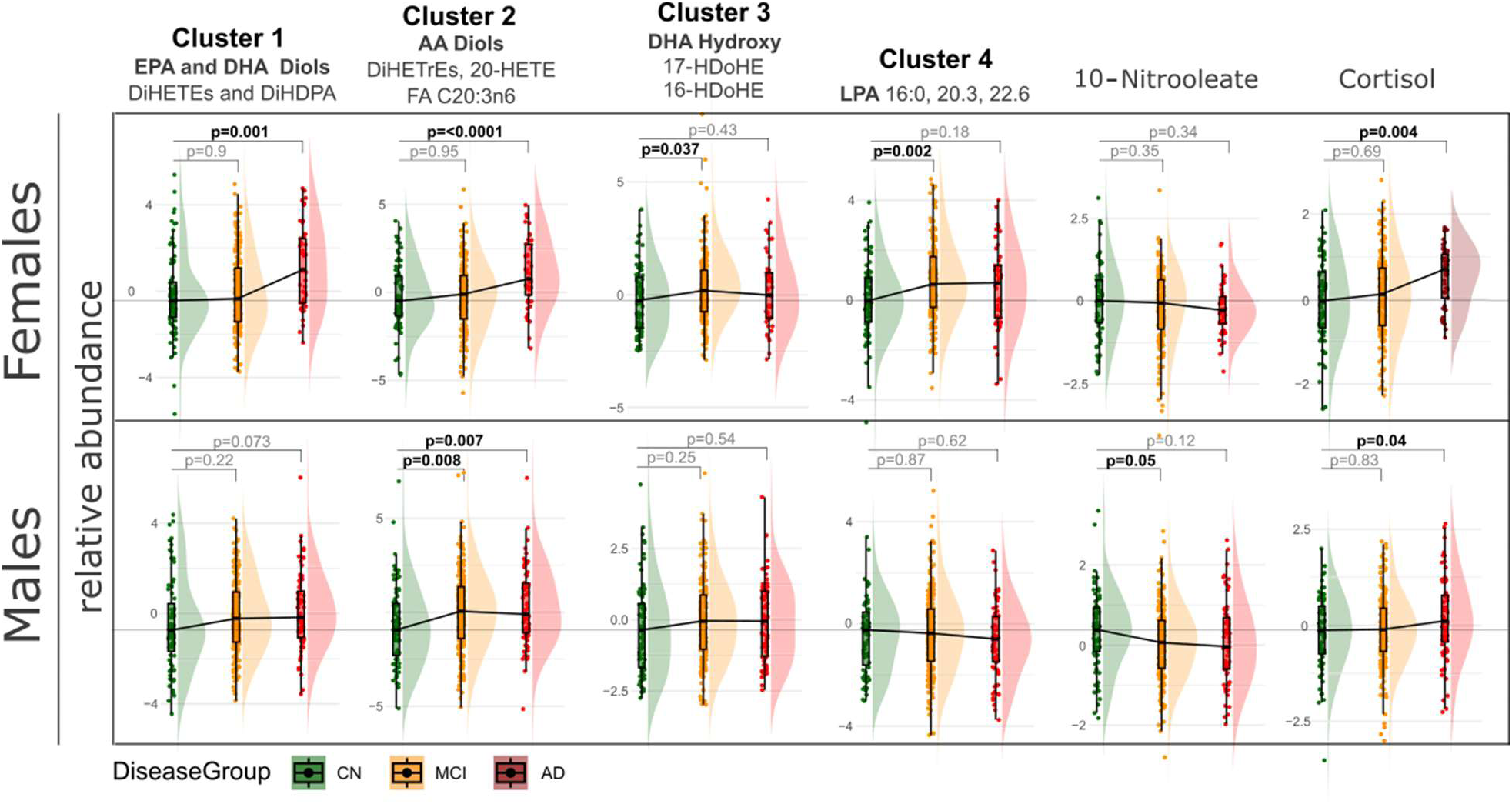
Soluble epoxide hydrolase metabolites are elevated along AD trajectory in a sex specific manner. Differences in means of lipid mediators between diagnosis groups, stratified by sex. To reduce data dimensionality and to facilitate interpretation and presentation, lipid mediators manifesting significant diagnosis group differences were converted into cluster components. Individual cluster members are defined in the cluster description, with the more detailed description of individual contribution to the cluster and differences in means between the treatment groups provided in the **Supplemental Table S1 and Supplemental Table S2**. Colors indicate diagnosis group: green – cognitively normal (CN; 83 females, 75 males) defined as healthy subjects who remind healthy after 4 years follow up; yellow - mild cognitively impaired (MCI; 179 females, 215 males); red – Alzheimer’s disease (AD; 53 females, 79 males). P values are derived from ANOVA with Dunnett post-test and indicate the differences from the CN.

In males, omega-3 diols differences did not reach significance (e.g., AD vs stable CN p =0.07). On the other hand, AA diols were higher in MCI and AD, compared to CN. This diagnosis group effect for both omega-3 and omega-6 vicinal diols was different from the one observed in females, with the diagnosis group x sex interaction p =0.078 (for cluster 1, omega-3 diols) and p =0.022 (for cluster 2, omega-6 diols) in the diagnosis group and sex factorial analysis. Additionally, 10-Nitrooleate was lower in MCI than CN and cortisol was higher in AD then CN.

### 2. Lipid mediators and APOE4 discriminate between stable MCI and MCI converters

Metabolic signatures associated with AD resilience, i.e., remaining MCI in comparison to MCI that progresses further (within 2 years) into AD, are poorly understood. Utilizing ADNI longitudinal information about patient diagnosis status, we defined the minimal number of factors (both clinical outcomes and lipid mediators, including bile acids) that can describe the difference between converters and non-converters in a sex-stratified analysis (**Figure 3**). In both sexes, combining APOE4 status with the plasma lipid mediator levels yielded the best discriminatory model: only APOE4 AUC = 64 for females and 62 for males; only lipid mediators AUC = 80 for females and 73 for males; APOE4 and lipid mediators AUC = 85 for females and 78 for males.

**Figure 3:**
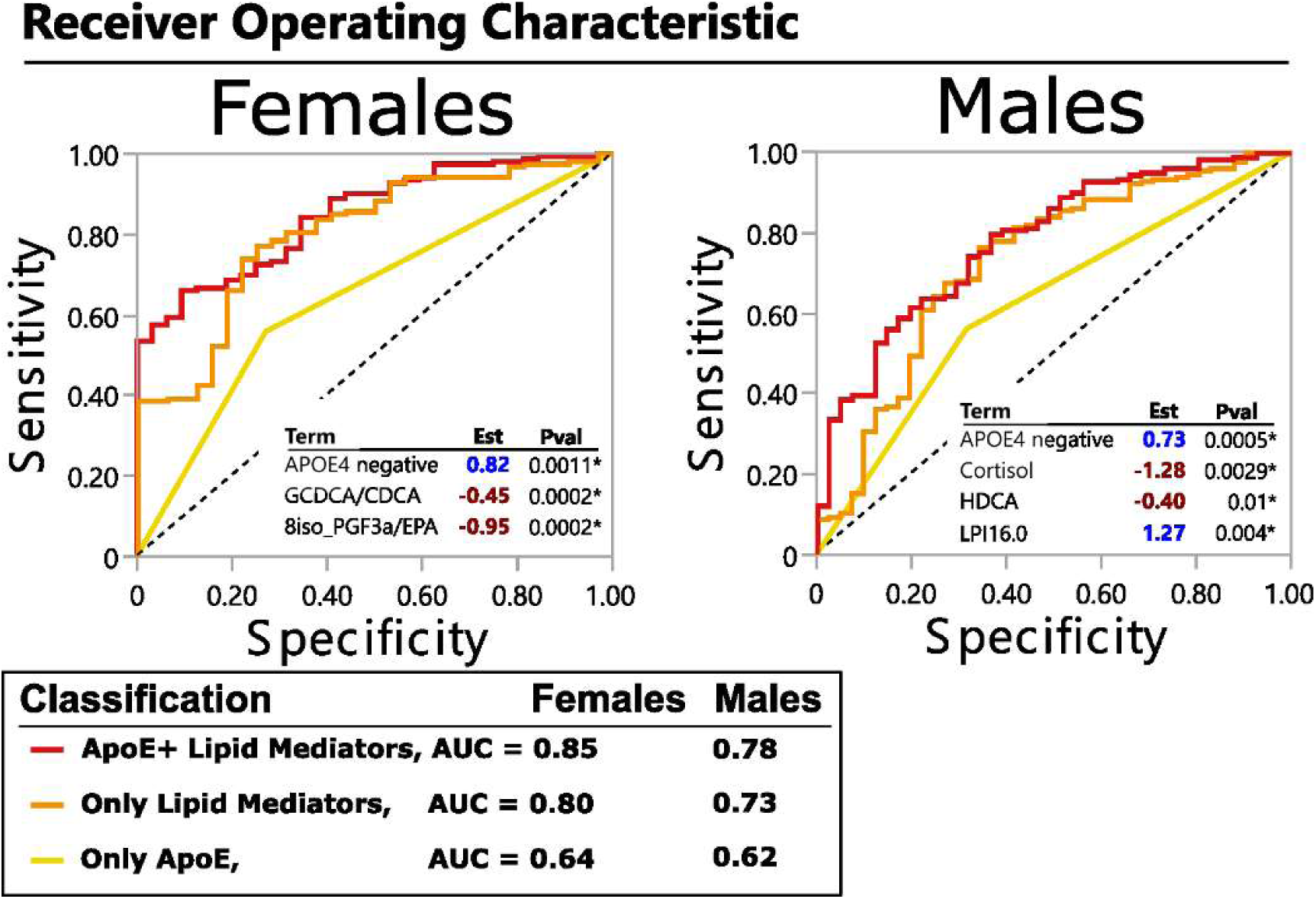
Lipid mediators and APOE genotype define AD resilient phenotype. Minimal model discriminating between stable MCI and MCI-AD converters (within 2 years), generated using stepwise logistic regression, stratified by sex. Biometrics (BMI, age,), presence of APOE4 allele and lipid mediators, including oxylipins, endocannabinoids and bile acids were used. The Receiver operation characteristics (ROC) curve is shown for the best model consisting of APOE4 information together with lipid mediators (red line) as well as for APOE4 alone (yellow line) and lipid mediators alone (orange line). The estimate and p values for each factor in the model are displayed in the ROC graph. The number of subjects: stable MCI = 153 females and 182 males; MCI converters = 32 females and 41 males.

In females, the likelihood of remaining MCI without converting to AD (AD resilience) was defined by the lack of an APOE4 allele, lower glycine conjugation of chenodeoxycholic acid (define as GCDCA/CDCA ratio) and lower levels of EPA-derived 8iso-prostaglanding F3 alpha (8-iso PGF3a). In males, AD resilience was defined by the lack of APOE4 allele, lower cortisol and hyodeoxycholic acid (HDCA) and higher levels of lysophosphatidylinostitol palmitate (i.e., LPI16.0). Further factorial analysis with group x sex interaction (model adjusted by APOE, age and BMI) confirm sex-specific effect for GCDCA/CDCA (p_interaction_=0.04), 8iso_PGF3a (p_interaction_=0.003) and cortisol (p_interaction_=0.04). The interaction terms for HDCA and LPI16:0 were not significant.

The derived minimal models had similar statistical power to discriminate converter vs non-converters whether the model was applied for the 1, 2 or 4 years follow up conversion criteria.

### 3. Markers of cognitive resilience among differential Tau and amyloid pathology, mediated by APOE genotypes

Next, we sought to define peripheral markers of memory function, measured by ADNI composite memory score(52) within the cohort. We are particularly interested in those markers in subjects manifesting high Tau and amyloid pathology, as indicators of cognitive resilience. Tau and amyloid pathology are measured by the ratio of total Tau to AB42 in CSF. In the ADNI GO/2 population, the Tau/AB42 ratio in CSF is influenced by APOE genotype, with e2 allele manifesting the least and e4 allele manifesting the highest level (**Supplemental Figure S2**). Each genotype manifested a range of memory scores, although e2e2 and e2e4 genotypes did not have sufficient number of participants for further analysis (n =3 and =14, respectively). To define the relationship between peripheral metabolism, Tau and amyloid pathology and memory, we applied stratification strategies for correlation analysis between plasma lipid mediators and the memory score. For correlation analysis, we applied a combination of random forest for features selection and stepwise linear regression to define a minimal model that explains the relation between memory and plasma lipid mediators (see materials and methods).

We identified APOE genotype stratification to yield significant models for associations of plasma metabolites and memory scores (**Figure 4**) in comparison to stratification by CSF Tau/AB42 (using previously published cutoff value (53)), which did not produce significant models. Each APOE genotype manifested unique sets of variables contributing to the model. Memory in APOE 2/3 was mainly defined by the age (negative association, p =0.0004) and 10-Nitrooleate (positive association, p =0.0005), in addition to lesser contribution of the ratio of GCDCA to GLCA (positive association, p =0.018) and LPG16:1 (negative association, p =0.015). Those 4 variables generated a memory model with the RSq =0.49 with the n =71. Memory in APOE3/3 genotype subjects was defined by age, the ratio of 19,20-DiHDPA/19,20-EpDPE, LPE22:5 (negative association, p =0.0001, 0.0079, 0.0028 respectively) and the years of education, LPI20:4 and the ratio of 5-HETE/AA (positive association, p =0.0001, 0.0013 and 0.015 respectively). Those 6 variables generated a memory model with the RSq =0.18 with the n =387. Memory in APOE3/4 genotype subjects was defined by age, 19,20-DiHDPA and LPE16:0 (negative association, p<0.0001, 0.005, and 0.019 respectively). Those 3 variables generated a memory model with the RSq=0.16 with the n=283. Memory in APOE4/4 genotype subjects was defined by an unknown (unidentified) oxylipin, lysophosphatidylinositol (LPI)16:1, 8HDoHE (positive association, p=0.0004, 0.0075 and 0.0096 respectively) and cortisol, leukotriene E4 (LTE4) and GDCA (negative association, p=0.0015, 0.0021, 0.0027 respectively). Those 6 variables generated a memory model with the RSq=0.46 with the n=78. The unknown (unidentified) oxylipin manifested mass transition of Resolvin E2 (m/z 333 -> m/z 115, the 115 fragment is for carboxyl group until 4’ carbon, as it breaks by 5’ hydroxyl group), but the chromatography retention time was different from the analytical standard. Additional correlative analysis shows a high correlation of this unknown oxylipin with EPA-derived vicinal diols (**Supplemental Figure S3**). Considering the importance of this metabolite in our predictive model and poor understanding of Resolvin biology, the full identification of this compound is of interest and is underway.

**Figure 4.**
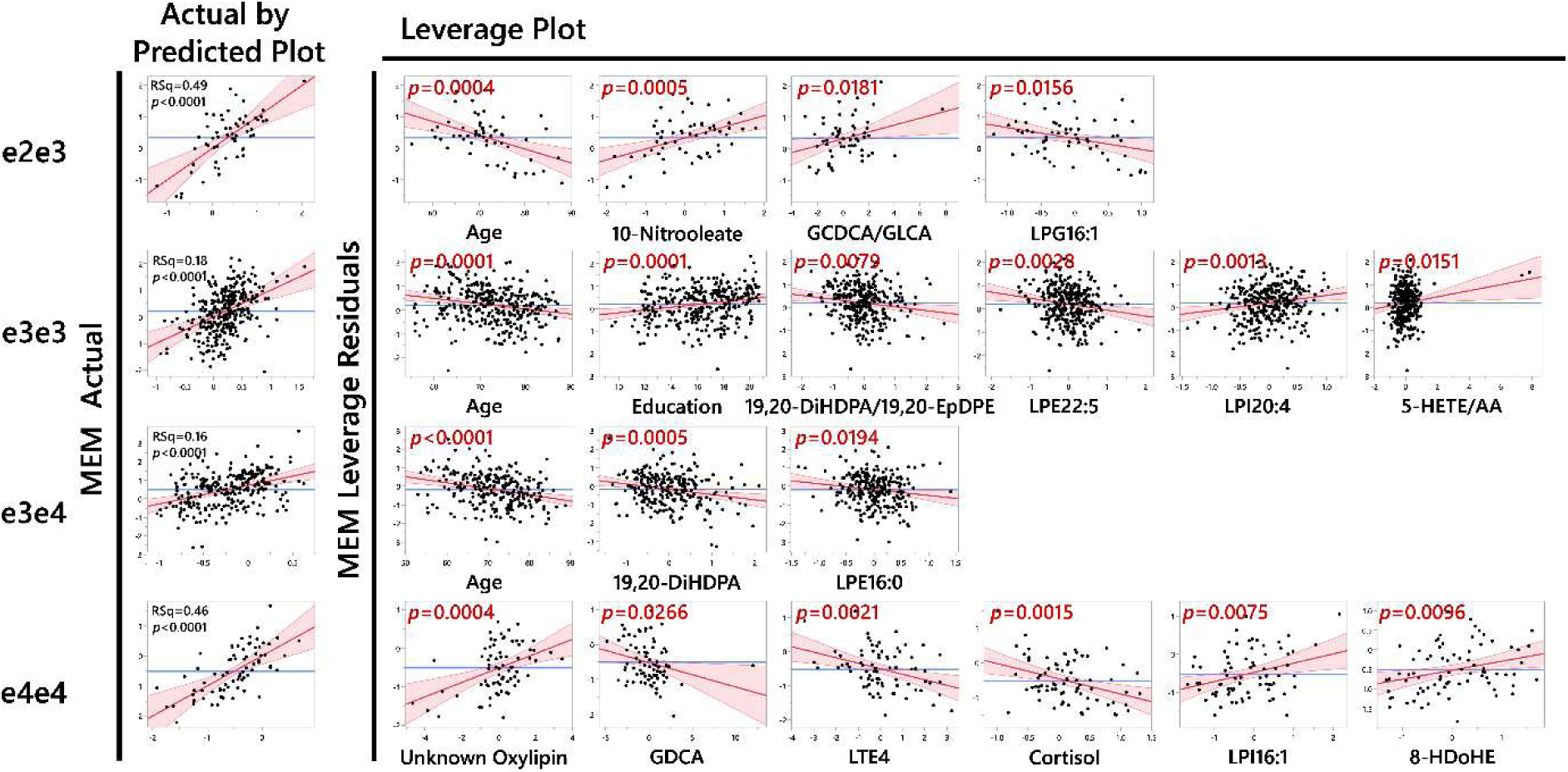
Lipid mediators reveal heterogeneity among APOE genotypes, an important AD risk factor. Multilinear model showing differential contribution of key peripheral metabolites and other factors such as age and education to memory scores (MEM) across APOE genotypes. The number of subjects: APOE2/3 = 60; APOE3/3 = 334; APOE3/4 = 254; APOE4/4 = 77.

To further demonstrate the APOE genotype specificity of those associations, we applied a factorial analysis for the key factors identified in the stepwise model (age, education, 10-Nitrooleate, 19,20-DiHDPA, unknown oxylipin, LTE4 and cortisol) to test the memory score interaction with APOE genotype. Following p_interactions_ were achieved with other key factors used as covariates in the model: age p_interaction_ = 0.06 for e4e4 vs e3e4; 0.35 for e4e4 vs e3e3 and 0.09 for e4e4 vs e2e3; education p_interaction_ = 0.67 for e3e3 vs e3e4; 0.45 for e3e3 vs e4e4 and 0.8 for e3e3 vs e2e3; 10-Nitrooleate p_interaction_ = 0.037 for e2e3 vs e3e3; 0.098 for e2e3 vs e3e4 and 0.74 for e2e3 vs e4e4; 19,20-DiHDPA p_interaction_ = 0.04 for e3e4 vs e4e4; 0.015 for e3e4 vs e3e3 and 0.05 for e3e4 vs e2e3; unknown oxylipin p_interaction_ = 0.12 for e4e4 vs e3e4; 0.07 for e4e4 vs e3e3 and 0.28 for e4e4 vs e2e3; LTE4 p_interaction_ = 0.05 for e4e4 vs e3e4; 0.0052 for e4e4 vs e3e3 and 0.0043 for e4e4 vs e2e3; cortisol p_interaction_ = 0.11 for e4e4 vs e3e4; 0.062 for e4e4 vs e3e3 and 0.17 for e4e4 vs e2e3;

Stepwise regression model identifies a minimal number of factors for predicted values and eliminates collinear variables. For example, 19,20-DiHDPA is highly correlated with other omega-3 diols, like EPA derived DiHETEs, and replacing 19,20-DiHDPA with closely correlated DiHETEs would yield similar predictive power. To illustrate the collinear relationship between measured lipid mediators and to facilitate minimal model interpretation, metabolite partial correlation network is presented in **Figure S3**.

## Discussion

Increased inflammatory responses have been reported in previous AD investigations but how these systems change over the progression of this disease have not been clearly delineated. Here we show a dysregulation of lipid mediators throughout AD progression and uncover disease heterogeneity influenced by sex and APOE genotype. We confirmed and extended our previous findings, from two independent cohorts, of increased soluble epoxide hydrolase activity during AD development. We also provide evidence implicating the APOE4 allele, bile acid metabolism, lipid peroxidation and stress response as sex-specific factors influencing the resistance of MCI subjects to progress to ADAPOE. Additionally, we showed APOE4/4 genotype-specific associations between plasma inflammatory markers and memory, identifying populations for precision medicine approach for potential therapeutic treatment.

Growing evidence points towards the sEH as a novel therapeutic target for AD (28, 54). Furthermore, plasma sEH metabolites (derived from AA, EPA and DHA) increase with AD pathology (53) and these metabolites are associated with cognition (i.e. perceptual speed) in healthy and MCI individuals (48). A strong connection of sEH-associated biochemistry with AD-affected vascular inflammation and energy metabolism in the central nervous system was also observed (55). Increased sEH expression in the AD-afflicted brain at both the transcript and protein levels are also seen (56). In rodent models of AD, AD-related pathologies can be altered by manipulating both sEH (28, 29) and CYP (30, 31) metabolism. Moreover, several genetic polymorphisms in the CYP/sEH pathway have consistently pointed towards reduced CYP-dependent epoxy fatty acid production and increased sEH activity or expression to be a risk for AD and increase AD-related pathologies (57–60), supporting causative effects of this critical inflammatory switch in the progression of this disease.

The current work provides additional evidence from a third independent AD cohort to support the dysregulation of she-dependent metabolism during AD development. Furthermore, the current study provides additional granularity regarding sex, APOE genotype and pre-AD stages. In particular, we saw an increase in EPA, DHA and AA-derived sEH products in females with AD, but not with MCI, compared to HC participants. In contrast, in males, we only saw an increase in AA-derived sEH metabolites, but they were present in subjects with MCI, suggesting that dysregulation of sEH in males may appear earlier in disease progression than in females. Similarly, differences in sEH metabolism have been reported to contribute to the sex difference in the vulnerability to other dementia risk factors, such as ischemic brain injury (61, 62), high glycemic diet (63–66) and high-fat diet (67, 68). At the molecular level, the possible involvement of sEH in sex-specific AD development includes a wide range fatty acid diol cellular functions (or depletion of fatty acids epoxide), e.g., ferroptosis-mediated neurodegeneration (69), exciting neurons (70), immune regulation (71, 72), and regulating mitochondrial respiration (73). For the epoxy fatty acids, their action is beneficial or detrimental depending on factors such as precursor fatty acids, concentration, targeting cells, and biological processes (71, 74). Together, this evidence suggests that reducing sEH activity may be a promising therapeutic approach in patients with both MCI and AD. These may include the use of sEH inhibitors currently under development (75, 76), but also include current therapies such as estrogen replacement therapy which can both down regulate sEH (77) and reduce AD risk in post-menopausal women (78).

Our results also support a sex-specific dysregulation of cortisol metabolism in AD, as seen by others (79, 80). Particularly we observed, increased cortisol concentrations in AD subjects when compared to CN in both sexes; increased levels in males MCI subjects that further convert to AD within 2 years, when compared to stable MCI phenotype; and negative association with memory in APOE 4/4 carriers. Cortisol is a stress response hormone that facilitates glucose dependent energy metabolism. This hormone crosses blood brain barrier via P-glycoprotein transporter (81) consistent with the strong correlation between plasma and CSF cortisol levels (53). Cortisol was previously implicated in neurodegeneration (82) and cognition (83) *via* regulation of the Hypothalamic-Pituitary-Adrenal (HPA) axis (84, 85) and was suggested by others as potential therapeutic target for AD (86). Sex-related differences were also reported, with plasma cortisol associations with brain volumes and memory performance being more pronounced in females than in males (87). We also show greater difference between CN and AD in females then in males. Our analysis here also adds to those findings by extending our understanding of cortisol and stress involvement in cognition and AD in specific subpopulations. In particular, we noted that males with MCI have elevated cortisol prior to conversion to AD, an effect not observed in females. This appears to contradict the previous findings suggesting a greater importance of this stress hormone in females with AD. Additionally in subjects with APOE 4/4 genotype, we show evidence that cortisol may impact memory performance, together with immune cells regulators. Our finding adds granularity to the known associations of stress hormones, cognition and AD development, identifying subpopulations for further potential therapeutic targeting of cortisol pathway in AD.

The current work also indicates the differential relationship between markers of immune cells activation and memory performance among various APOE genotype carriers. Notably in the presence of the APOE 4/4 genotype, memory was negatively associated with the pro-inflammatory cysteinyl leukotriene metabolism reported by LTE4, generated *via* 5-LOX-dependent metabolism. LTE4 is a monocyte activator with vasoactive properties, activating vascular permeability and edema (88), and its levels were associated with cardiovascular disorders (89, 90). Previously, 5-LOX expression was found to be upregulated in the peripheral blood mononuclear cells from AD patients (91, 92). Furthermore, a gain of function 5-LOX polymorphism was also associated with risk of AD (93), indicating a potential causative relationship of the dysregulation of this pathway to AD. Meanwhile in rodent models of AD upregulation of hippocampal and cortex 5-LOX expression was associated with elevated γ-secretase activity and Aβ peptide formation (92, 94–97). APOE genotypes-specific oxylipin profile in postmortem human dorsolateral prefrontal cortex found that 5-LOX-derived EPA oxylipins LXA5 and LXB5 in brain were negatively associated with cognitive performance, and the association was stronger among APOE4 carriers compared to APOE3 carriers (98). This research also found that the levels of a 5-LOX product, LXA4 was negatively associated with sEH activity. On the basis of this, our results here further indicate that the dynamics between APOE genotypes, 5-LOX, sEH, fatty acid precursors and cognition can be captured in the blood lipidome, which can be monitored less invasively and more routinely for longitudinal cohorts.

The functional interaction between APOE, 5-LOX and sEH can be supported by various mechanisms: for example, the higher level of APOE4 carriers in activation of Ca^2+^ influx and thus Ca^2+^-dependent cPLA2 and 5-LOX compared to APOE2 or E3 (98, 99); for another, the presence of prone-to-aggregation lipid-poor APOE4 protein can affect the lipid efflux functions of ABCA1(100) and change the lipid composition of cells (101), which serve as substrates to release free fatty acids subject to 5-LOX and sEH metabolism (102, 103). A possibility also remains that APOE4 leads to the altered ability of trafficking inflammation-regulating lipid mediators between lipoprotein and lipoprotein-metabolizing cells, which includes microglia, a critical player in AD pathogenesis (102). Further investigation is therefore warranted to clarify if the combination of altering 5-LOX metabolism and fatty acid composition together can be a specific modifiable target for APOE4 carriers to improve cognition.

## Limitations

This study is not big enough to explore APOE genotypes and sex combined stratification. ADNI 1 and GO/2 cohorts do not represent full demographics of US and world population. This shortcoming is addressed in ADNI 3 cohort. Longitudinal data are needed to further connect dysregulation in lipid mediators, sEH among others and disease progression, define subgroups to identify population for targeted interventions.

## Acknowledgement

The Alzheimer’s Disease Metabolomics Consortium (ADMC) is funded wholly or in part by the following National Institute on Aging grants and supplements, which are components of the Accelerating Medicines Partnership for AD (AMP-AD) and/or Molecular Mechanisms of the Vascular Etiology of AD (M2OVE-AD): NIA R01AG046171, RF1AG051550, RF1AG057452, R01AG059093, RF1AG058942, U01AG061359, U19AG063744, 3U19AG063744-04S1, 1R01AG081322, U01AG088562, and FNIH: #DAOU16AMPA awarded to Dr. Rima Kaddurah-Daouk at Duke University in partnership with a large number of academic institutions. Chunyuan Yin acknowledges support from the China Scholarship Council fellowship (No. 202006550003). TH acknowledges funding for the Exposome-Scan infrastructure by the Dutch Research Council (project number 175.2019.032).

## Data availability statement

All metabolomics data is publicly available at https://ida.loni.usc.edu/login.jsp

