## Supplemental Figure S1 for "Metabolic Alteration in Oxylipins and Endocannabinoids Point to an Important Role for Soluble Epoxide Hydrolase and Inflammation in Alzheimer’s Disease - Finding from Alzheimer’s Disease Neuroimaging Initiative"

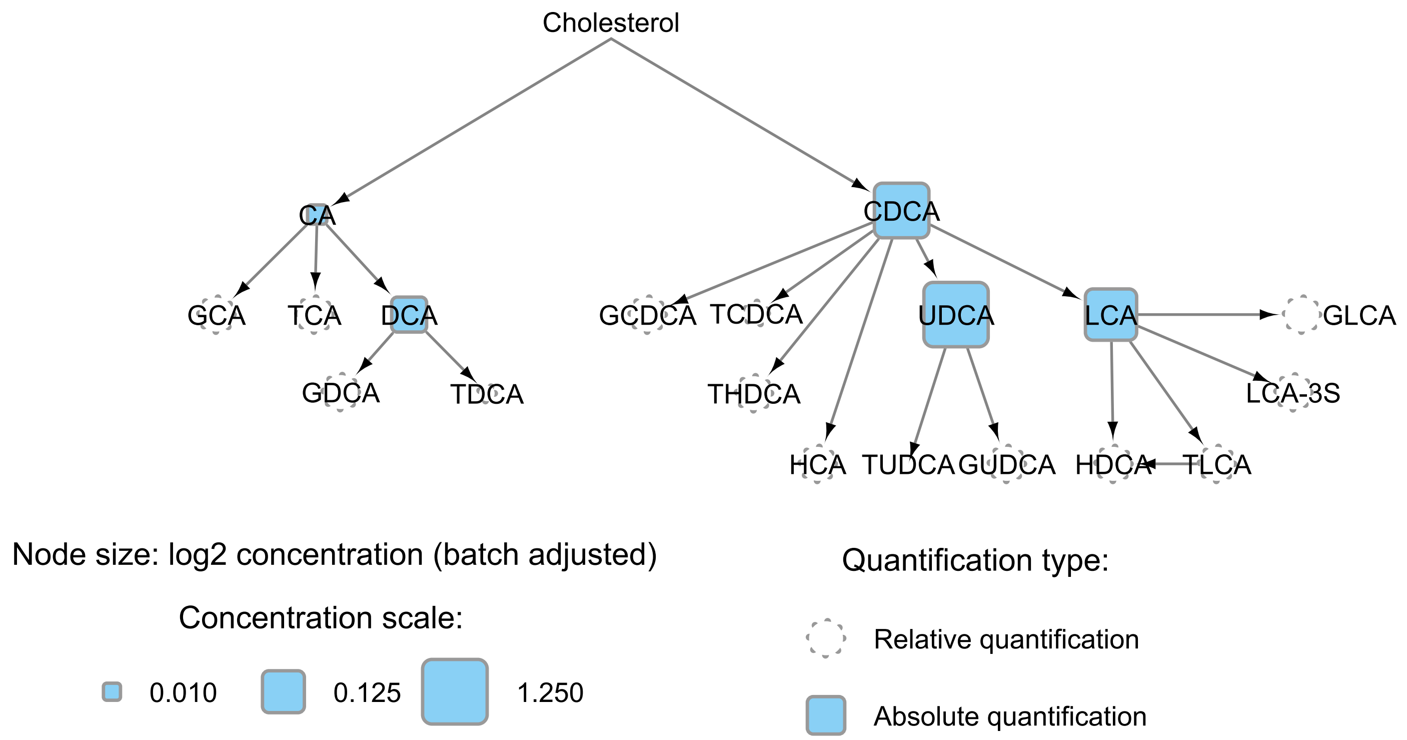


**Supplemental Figure S1. Quantified bile acids projected onto their metabolic pathway with node sizes indicating their average concentration in samples.**
