## Supplemental Figure S2 for "Metabolic Alteration in Oxylipins and Endocannabinoids Point to an Important Role for Soluble Epoxide Hydrolase and Inflammation in Alzheimer’s Disease - Finding from Alzheimer’s Disease Neuroimaging Initiative"

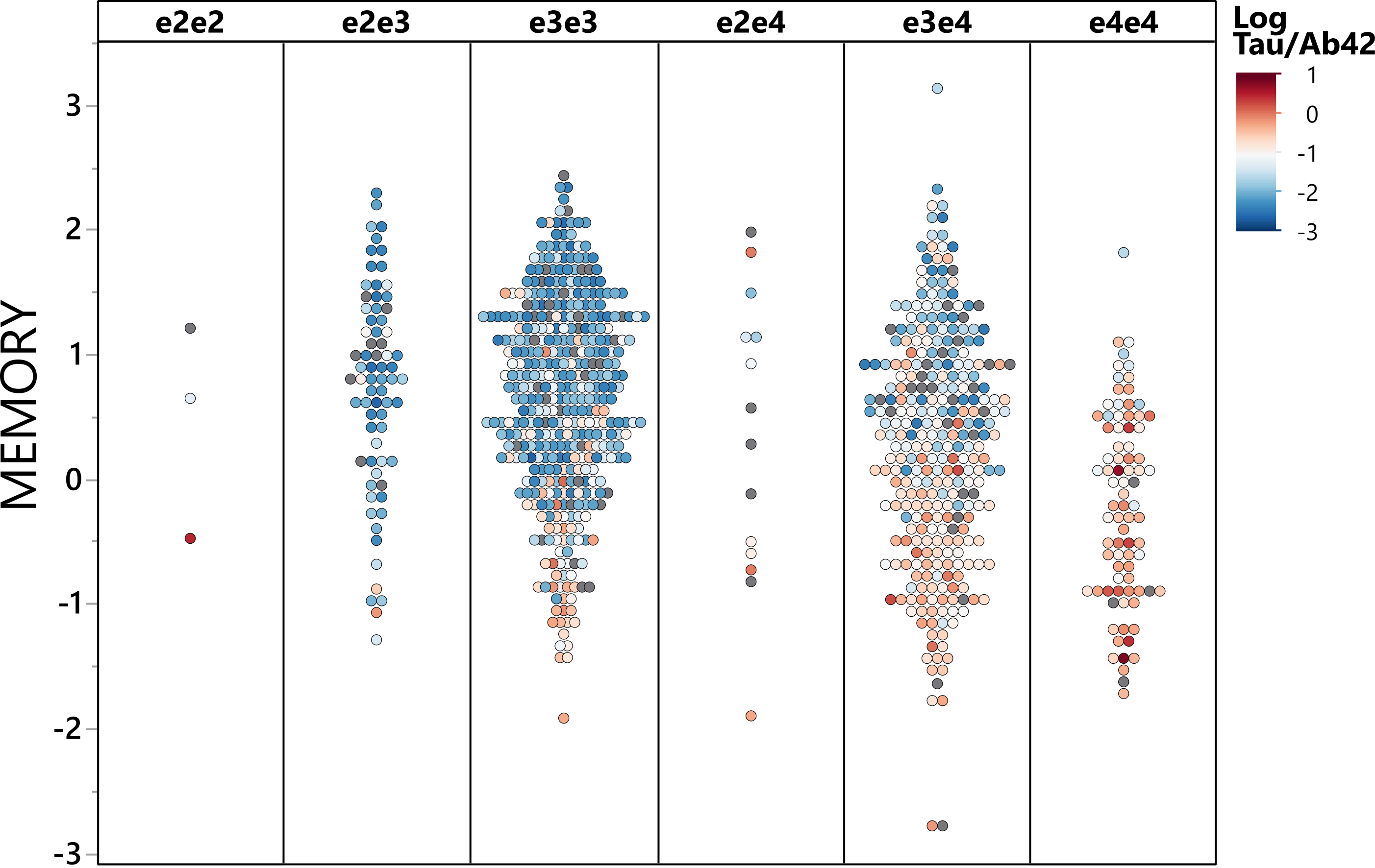


**Supplemental Figure S2. Relation between Tau and Ab42 CSF biomarkers, the memory score and APOE genotype in selected ADNI2 subjects.**
