## Supplementary figures and images for "Metabolic Alteration in Oxylipins and Endocannabinoids Point to an Important Role for Soluble Epoxide Hydrolase and Inflammation in Alzheimer’s Disease - Finding from Alzheimer’s Disease Neuroimaging Initiative"

### Supplemental Figure S3

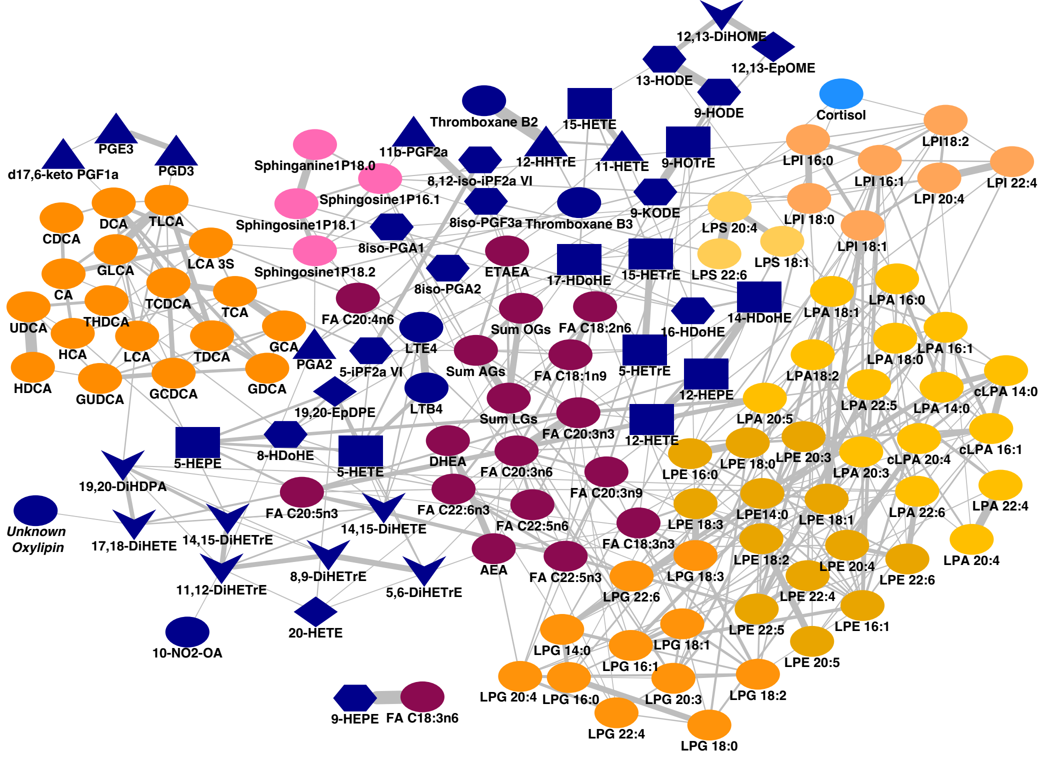
