## Supplemental Table S1 for "Metabolic Alteration in Oxylipins and Endocannabinoids Point to an Important Role for Soluble Epoxide Hydrolase and Inflammation in Alzheimer’s Disease - Finding from Alzheimer’s Disease Neuroimaging Initiative"

Supplemental Table S1. cluster members as well as their correlations within the cluster

| Cluster | Metabolites | RSquare with Own Cluster | RSquare with Next Closest | 1-RSquare Ratio |
|---|---|---|---|---|
| 1 | sum_DiHETE | 0.95 | 0.42 | 0.09 |
| 1 | 17_18_DiHETE | 0.91 | 0.39 | 0.15 |
| 1 | 19_20_DiHDPA | 0.75 | 0.32 | 0.36 |
| 1 | 14_15_DiHETE | 0.75 | 0.40 | 0.41 |
| 2 | GLCA/CA | 0.83 | 0.30 | 0.24 |
| 2 | GDCA/CA | 0.83 | 0.36 | 0.27 |
| 2 | HDCA/CA | 0.76 | 0.02 | 0.25 |
| 2 | UDCA/CA | 0.74 | 0.02 | 0.26 |
| 2 | GCA/CA | 0.61 | 0.12 | 0.44 |
| 2 | LCA_3S/CA | 0.58 | 0.16 | 0.50 |
| 3 | sum_DiHETrE | 0.91 | 0.19 | 0.11 |
| 3 | 11_12_DiHETrE | 0.87 | 0.20 | 0.16 |
| 3 | 14_15_DiHETrE | 0.84 | 0.18 | 0.19 |
| 3 | 8_9_DiHETrE | 0.78 | 0.16 | 0.26 |
| 3 | 20_HETE | 0.61 | 0.12 | 0.45 |
| 3 | FA20.3_w6 | 0.35 | 0.12 | 0.74 |
| 4 | LPA16.0 | 0.84 | 0.13 | 0.19 |
| 4 | LPA20.3 | 0.78 | 0.20 | 0.27 |
| 4 | LPA22.6 | 0.67 | 0.40 | 0.54 |
| 4 | LPI16.0 | 0.30 | 0.03 | 0.73 |
| 5 | 19_20_DiHDPA/DHA | 0.70 | 0.07 | 0.33 |
| 5 | 17_18_DiHETE/EPA | 0.66 | 0.15 | 0.40 |
| 5 | 14_15_DiHETE/EPA | 0.58 | 0.14 | 0.49 |
| 5 | 19_20_DiHDPA/19_20_EpDPE | 0.46 | 0.12 | 0.62 |
| 5 | 14,15-DiHETrE/AA | 0.30 | 0.07 | 0.75 |
| 6 | 1_AG_2_AG | 0.88 | 0.09 | 0.13 |
| 6 | 1_LG_2_LG | 0.88 | 0.03 | 0.13 |
| 7 | GDCA | 0.83 | 0.13 | 0.20 |
| 7 | GLCA | 0.83 | 0.17 | 0.21 |
| 8 | sum_HDoHE | 0.93 | 0.26 | 0.09 |
| 8 | 17_HDoHE | 0.69 | 0.26 | 0.42 |
| 8 | 16_HDoHE | 0.62 | 0.15 | 0.45 |
| 9 | 5_HEPE | 0.85 | 0.37 | 0.23 |
| 9 | 8_HDoHE | 0.79 | 0.40 | 0.34 |
| 9 | 19_20_EpDPE | 0.70 | 0.22 | 0.38 |
| 9 | PGD3/EPA | 0.52 | 0.29 | 0.69 |
| 10 | 10_NO2_OA | 1.00 | 0.00 | 0.00 |
| 11 | Cortisol | 1.00 | 0.03 | 0.00 |
