## Supplemental Table S2 for "Metabolic Alteration in Oxylipins and Endocannabinoids Point to an Important Role for Soluble Epoxide Hydrolase and Inflammation in Alzheimer’s Disease - Finding from Alzheimer’s Disease Neuroimaging Initiative"

| Cluster | Gender | Anova p value | DunnettTest- MCI to HC | DunnettTest- AD to HC |
|---|---|---|---|---|
| Cluster 1 | male | 0.12 | 0.22 | 0.073 |
| Cluster 1 | female | 0.00013 | 0.92 | 0.00028 |
| Cluster 2 | male | 0.00096 | 0.04 | 0.00038 |
| Cluster 2 | female | 0.68 | 0.66 | 0.67 |
| Cluster 3 | male | 0.0056 | 0.0084 | 0.0067 |
| Cluster 3 | female | 0.00018 | 0.95 | 0.00042 |
| Cluster 4 | male | 0.38 | 0.87 | 0.63 |
| Cluster 4 | female | 0.0039 | 0.0017 | 0.18 |
| Cluster 5 | male | 0.0051 | 0.95 | 0.011 |
| Cluster 5 | female | 0.0000027 | 0.61 | 0.00018 |
| Cluster 6 | male | 0.0063 | 0.44 | 0.13 |
| Cluster 6 | female | 0.48 | 0.39 | 0.92 |
| Cluster 7 | male | 0.15 | 0.74 | 0.11 |
| Cluster 7 | female | 0.45 | 0.88 | 0.35 |
| Cluster 8 | male | 0.36 | 0.26 | 0.54 |
| Cluster 8 | female | 0.065 | 0.037 | 0.43 |
| Cluster 9 | male | 0.24 | 0.34 | 0.97 |
| Cluster 9 | female | 0.16 | 0.13 | 0.22 |
| Cluster 10 | male | 0.077 | 0.053 | 0.13 |
| Cluster 10 | female | 0.35 | 0.35 | 0.34 |
| Cluster 11 | male | 0.038 | 0.83 | 0.038 |
| Cluster 11 | female | 0.006 | 0.69 | 0.0045 |
