## Supplemental Table S3 for "Metabolic Alteration in Oxylipins and Endocannabinoids Point to an Important Role for Soluble Epoxide Hydrolase and Inflammation in Alzheimer’s Disease - Finding from Alzheimer’s Disease Neuroimaging Initiative"

Supplemental Table S3. P values from ANOVA analysis diagnosis group vs metabolites and informative ratios and summations

| Metabolite | Group | Group * Sex Int | APOEGrp (0 = e4 negative; 1 = e4 positive) | BMI | Education | Gender | Age |
|---|---|---|---|---|---|---|---|
| 19_20_DiHDPA/19_20_EpDPE | **<.0001** | 0.13 | 0.51 | 0.0011 | 0.82 | 0.97 | 0.33 |
| 14_15_DiHETrE | **<.0001** | **0.048** | 0.23 | 0.01 | 0.19 | 0.57 | 0.38 |
| 19_20_DiHDPA | **<.0001** | **0.0016** | 0.9 | <.0001 | 0.032 | 0.0001 | 0.01 |
| 11_12_DiHETrE | **<.0001** | 0.055 | 0.12 | 0.0003 | 0.44 | 0.98 | 0.2 |
| sum_DiHETrE | **<.0001** | **0.023** | 0.15 | 0.001 | 0.86 | 0.53 | 0.15 |
| 8_9_DiHETrE | **0.0001** | **0.049** | 0.33 | 0.06 | 0.92 | 0.14 | 0.15 |
| 20_HETE | **0.0002** | 0.063 | 0.059 | 0.0074 | 0.7 | 0.11 | 0.11 |
| 17_18_DiHETE/EPA | **0.0002** | 0.82 | 0.73 | 0.0049 | 0.96 | 0.015 | 0.62 |
| 19_20_DiHDPA/DHA | **0.0002** | 0.67 | 0.88 | 0.0007 | 0.86 | 0.061 | 0.45 |
| Cortisol | **0.001** | 0.43 | 0.77 | 0.048 | 0.47 | 0.01 | 0.27 |
| 14_15_DiHETE/EPA | **0.0011** | 0.87 | 0.3 | 0.039 | 0.31 | 0.0004 | 0.084 |
| GDCA/CA | **0.0018** | 0.36 | 0.54 | 0.47 | 0.056 | 0.21 | 0.12 |
| FA20.3_w6 | **0.002** | 0.11 | 0.52 | 0.22 | 0.63 | <.0001 | 0.96 |
| GLCA/CA | **0.0038** | 0.58 | 0.83 | 0.2 | 0.2 | 0.24 | 0.28 |
| UDCA/CA | **0.0042** | 0.36 | 0.66 | 0.23 | 0.46 | 0.61 | 0.16 |
| HDCA/CA | **0.0044** | 0.29 | 0.97 | 0.2 | 0.3 | 0.17 | 0.11 |
| 17_18_DiHETE | **0.0054** | 0.22 | 0.87 | <.0001 | 0.0049 | 0.0007 | 0.057 |
| GDCA | **0.0055** | 0.89 | 0.067 | 0.17 | 0.034 | 0.97 | 0.99 |
| GLCA | **0.0065** | 0.39 | 0.48 | 0.68 | 0.16 | 0.72 | 0.63 |
| LCA_3S/CA | **0.0066** | 0.61 | 0.47 | 0.18 | 0.34 | 0.92 | 0.62 |
| GCDCA/CA | **0.0078** | 0.86 | 0.86 | 0.16 | 0.15 | 0.96 | 0.24 |
| sum_DiHETE | **0.0094** | 0.18 | 0.96 | <.0001 | 0.0057 | 0.0016 | 0.073 |
| 1_AG_2_AG | **0.011** | 0.11 | 0.27 | 0.8 | 0.28 | 0.35 | 0.56 |
| LCA | **0.012** | 0.92 | 0.19 | 0.11 | 0.74 | 0.8 | 0.17 |
| LPA16.0 | **0.021** | 0.15 | 0.92 | 0.7 | 0.95 | <.0001 | 0.02 |
| LCA/CA | **0.022** | 0.074 | 0.97 | 0.31 | 0.55 | 0.15 | 0.22 |
| DCA/CA | **0.023** | **0.017** | 0.88 | 0.87 | 0.27 | 0.24 | 0.026 |
| 19_20_EpDPE | **0.028** | 0.4 | 0.76 | 0.017 | 0.016 | 0.0006 | 0.21 |
| 14_15_DiHETE | **0.028** | 0.35 | 0.91 | <.0001 | 0.022 | 0.041 | 0.31 |
| GUDCA/CA | **0.029** | 0.56 | 0.6 | 0.18 | 0.073 | 0.13 | 0.24 |
| LPA22.6 | **0.031** | 0.32 | 0.81 | 0.0036 | 0.02 | <.0001 | 0.0032 |
| Sphingosine1P18.1 | **0.033** | 0.52 | 0.96 | 0.18 | 0.019 | 0.033 | 0.046 |
| PGD3/EPA | **0.034** | 0.76 | 0.89 | 0.071 | 0.34 | <.0001 | 0.61 |
| UDCA | **0.034** | 0.75 | 0.044 | 0.44 | 0.54 | 0.18 | 0.14 |
| TLCA/CA | **0.036** | 0.4 | 0.82 | 0.24 | 0.47 | 0.051 | 0.3 |
| GCA/CA | **0.036** | 0.89 | 0.74 | 0.53 | 0.31 | 0.24 | 0.13 |
| THDCA/CA | **0.038** | 0.17 | 0.92 | 0.28 | 0.54 | 0.61 | 0.0062 |
| 8_HDoHE | **0.042** | 0.31 | 0.72 | 0.0099 | 0.0002 | <.0001 | 0.0046 |

| | | | | | | | |
|---|---|---|---|---|---|---|---|
| HDCA | **0.042** | 0.44 | 0.18 | 0.7 | 0.2 | 0.44 | 0.18 |
| TLCA | **0.043** | 0.86 | 0.18 | 0.8 | 0.55 | 0.13 | 0.29 |
| GLCA/CDCA | **0.045** | 0.91 | 0.25 | 0.2 | 0.13 | 0.14 | 0.8 |
| TDCA/CA | **0.046** | 0.36 | 0.89 | 0.45 | 0.47 | 0.052 | 0.13 |
| 1_LG_2_LG | 0.051 | 0.11 | 0.049 | 0.43 | 0.011 | 0.93 | 0.4 |
| 5_HETrE/DGLA | 0.055 | 0.36 | 0.22 | 0.82 | 0.34 | 0.67 | 0.58 |
| 8_12_iso_iPF2a_VI/AA | 0.059 | 0.069 | 0.4 | 0.0009 | 0.66 | 0.51 | 0.2 |
| GDCA/DCA | 0.059 | 0.65 | 0.43 | 0.31 | 0.11 | 0.82 | 0.74 |
| FA22.6_w3 | 0.064 | **0.022** | 0.84 | 0.0069 | 0.006 | <.0001 | 0.079 |
| 14,15-DiHETrE/AA | 0.067 | 0.55 | 0.84 | 0.11 | 0.66 | 0.0013 | 0.36 |
| 15-HETrE/DGLA | 0.067 | 0.084 | 0.35 | 0.58 | 0.49 | 0.055 | 0.46 |
| CDCA/CA | 0.07 | 0.3 | 0.3 | 0.95 | 0.94 | 0.86 | 0.033 |
| THDCA | 0.073 | 1 | 0.15 | 0.34 | 0.56 | 0.35 | 0.11 |
| LPI16.0 | 0.079 | **0.024** | 0.9 | 0.71 | 0.61 | 0.0007 | 0.66 |
| LPA22.5 | 0.081 | 0.52 | 0.86 | 0.025 | 0.7 | <.0001 | 0.59 |
| LPA20.5 | 0.082 | 0.43 | 0.33 | 0.044 | 0.0085 | <.0001 | 0.011 |
| 19_20_EpDPE/DHA | 0.082 | 0.3 | 0.43 | 0.81 | 0.95 | 0.15 | 0.78 |
| FA20.4_w6 | 0.084 | 0.33 | 0.35 | 0.85 | 0.83 | 0.0009 | 0.84 |
| LPA20.3 | 0.084 | 0.74 | 0.85 | 0.23 | 0.99 | <.0001 | 0.74 |
| Sphinganine1P18.0 | 0.087 | 0.67 | 0.65 | 0.34 | 0.24 | 0.0002 | 0.3 |
| DHEA | 0.087 | 0.068 | 0.073 | 0.86 | 0.017 | 0.077 | 0.015 |
| 5_HEPE | 0.088 | 0.5 | 0.6 | 0.13 | 0.087 | <.0001 | 0.09 |
| LPA20.4 | 0.088 | 0.95 | 0.86 | 0.4 | 0.29 | 0.0095 | 0.36 |
| 16_HDoHE | 0.099 | 0.99 | 0.89 | 0.1 | 1 | 0.28 | 0.23 |
| TCDCA/CA | 0.1 | 0.85 | 0.95 | 0.21 | 0.25 | 0.057 | 0.05 |
| 16-HDoHE/DHA | 0.1 | 0.97 | 0.85 | 0.26 | 0.41 | 0.75 | 0.15 |
| TDCA | 0.1 | 0.87 | 0.24 | 0.41 | 0.71 | 0.18 | 0.96 |
| GCDCA | 0.1 | **0.046** | 0.19 | 0.82 | 0.17 | 0.098 | 0.37 |
| LPA16.1 | 0.1 | 0.19 | 0.86 | 0.017 | 0.47 | <.0001 | 0.0083 |
| 11_12_DiHETrE/14_15_DiHETrE | 0.11 | 0.87 | 0.55 | 0.043 | 0.55 | 0.76 | 0.41 |
| 14_15_DiHETrE/11_12_DiHETrE | 0.11 | 0.87 | 0.55 | 0.043 | 0.55 | 0.76 | 0.41 |
| 5_6_DiHETrE | 0.11 | **0.048** | 0.18 | 0.025 | 0.25 | 0.37 | 0.063 |
| LPE22.6 | 0.12 | 0.053 | 0.92 | <.0001 | 0.013 | <.0001 | 0.001 |
| HCA/HDCA | 0.12 | 0.53 | 0.98 | 0.61 | 0.12 | 0.39 | 0.96 |
| LPS18.1 | 0.12 | 0.91 | 0.46 | 0.14 | 0.66 | 0.22 | 0.35 |
| GCDCA/CDCA | 0.12 | 0.39 | 0.47 | 0.21 | 0.16 | 0.87 | 0.54 |
| LPA18.0 | 0.12 | 0.37 | 0.69 | 0.34 | 0.77 | <.0001 | 0.88 |
| 12_HETE | 0.14 | 0.88 | 0.5 | 0.34 | 0.97 | 0.048 | 0.29 |
| (GCA+TCA)/CA | 0.15 | 0.34 | 0.56 | 0.82 | 0.54 | 0.065 | 0.079 |
| 10_NO2_OA | 0.15 | 0.14 | 0.34 | 0.0012 | 0.77 | 0.0003 | 0.044 |
| LPA22.4 | 0.15 | 0.7 | 0.96 | 0.76 | 0.066 | 0.051 | 0.25 |
| PGD3 | 0.15 | 0.95 | 0.41 | 0.24 | 0.045 | 0.75 | 0.23 |
| TCA/CA | 0.16 | 0.33 | 0.56 | 0.83 | 0.55 | 0.06 | 0.079 |
| GCA/GDCA | 0.16 | 0.29 | 0.88 | 0.64 | 0.13 | 0.8 | 0.31 |
| FA18.1_w9 | 0.16 | **0.034** | 0.84 | 0.3 | 0.69 | <.0001 | 0.066 |

| | | | | | | | |
|---|---|---|---|---|---|---|---|
| LPI16.1 | 0.16 | 0.38 | 0.92 | 0.0005 | 0.81 | <.0001 | 0.92 |
| 8-HDoHE/DHA | 0.17 | **0.035** | 0.58 | 0.75 | 0.007 | 0.65 | 0.0056 |
| LPS20.4 | 0.17 | 0.57 | 0.54 | 0.22 | 0.22 | 0.28 | 0.17 |
| sum_HETE | 0.17 | 0.58 | 0.17 | 0.34 | 0.38 | 0.0011 | 0.57 |
| 12_HEPE | 0.17 | 0.82 | 0.69 | 0.011 | 0.4 | 0.0015 | 0.81 |
| FA20.5_w3 | 0.17 | 0.82 | 0.55 | 0.0035 | 0.005 | <.0001 | 0.058 |
| 12-HEPE/EPA | 0.17 | 0.75 | 0.62 | 0.22 | 0.46 | 0.22 | 0.57 |
| 14-HDoHE/DHA | 0.17 | 0.81 | 0.81 | 0.2 | 0.55 | 0.29 | 0.69 |
| HCA/CA | 0.18 | 0.66 | 0.78 | 0.054 | 0.75 | 0.73 | 0.059 |
| TLCA/CDCA | 0.18 | 0.91 | 0.52 | 0.32 | 0.36 | 0.0046 | 0.56 |
| cLPA20.4 | 0.18 | 0.68 | 0.77 | 0.47 | 0.097 | 0.14 | 0.0042 |
| 8iso_PGA1/AA | 0.19 | 0.85 | 0.14 | 0.46 | 0.84 | 0.011 | 0.81 |
| 11b_PGF2a | 0.19 | 0.84 | 0.26 | 0.83 | <.0001 | 0.25 | 0.0094 |
| LCA_3S | 0.19 | 0.66 | 0.099 | 0.56 | 0.36 | 0.022 | 0.012 |
| 8iso_PGF3a/EPA | 0.21 | 0.63 | 0.42 | 0.17 | 0.65 | <.0001 | 0.093 |
| 11b_PGF2a/AA | 0.21 | 0.87 | 0.18 | 0.93 | 0.0014 | 0.36 | 0.091 |
| 17_HDoHE | 0.21 | 0.89 | 0.42 | 0.07 | 0.5 | 0.91 | 0.35 |
| 12-HETE/AA | 0.21 | 0.95 | 0.76 | 0.27 | 0.85 | 0.42 | 0.22 |
| 14_HDoHE | 0.22 | 0.87 | 0.78 | 0.026 | 0.63 | 0.024 | 0.97 |
| TCDCA | 0.22 | 0.14 | 0.23 | 0.8 | 0.21 | 0.53 | 0.95 |
| FA20.3_w3 | 0.22 | **0.049** | 0.45 | 0.14 | 0.22 | <.0001 | 0.62 |
| ETAEA | 0.23 | 0.1 | 0.8 | 0.0086 | 0.6 | 0.44 | 0.13 |
| FA22.5_w3 | 0.23 | 0.094 | 0.58 | 0.025 | 0.12 | <.0001 | 0.17 |
| LPA18.1 | 0.23 | 0.075 | 0.46 | 0.06 | 0.37 | <.0001 | 0.22 |
| CA | 0.24 | 0.054 | 0.46 | 0.058 | 0.64 | 0.12 | 0.015 |
| LPG16.1 | 0.24 | 0.32 | 0.68 | <.0001 | 0.6 | <.0001 | 0.75 |
| DCA | 0.24 | 0.58 | 0.23 | 0.0012 | 0.21 | 0.78 | 0.98 |
| 8_12_iso_iPF2a_VI | 0.24 | **0.045** | 0.91 | <.0001 | 0.2 | <.0001 | 0.03 |
| UDCA/CDCA | 0.25 | 0.9 | 0.59 | 0.33 | 0.41 | 0.18 | 0.49 |
| LPE20.4 | 0.25 | 0.52 | 0.66 | 0.0075 | 0.053 | 0.33 | 0.38 |
| 9_HEPE | 0.26 | **0.014** | 0.75 | 0.39 | 0.096 | <.0001 | 0.1 |
| GCA | 0.26 | 0.22 | 0.2 | 0.17 | 0.44 | 0.61 | 0.37 |
| 8iso_PGA2/AA | 0.27 | 0.4 | 0.081 | 0.18 | 0.2 | <.0001 | 0.2 |
| 11,12-DiHETrE/AA | 0.27 | 0.62 | 0.98 | 0.0094 | 0.81 | 0.0005 | 0.34 |
| LCA/CDCA | 0.29 | 0.54 | 0.26 | 0.65 | 0.51 | 0.091 | 0.52 |
| sum_HDoHE | 0.29 | 0.81 | 0.74 | 0.0053 | 0.42 | 0.015 | 0.41 |
| PGA2 | 0.29 | 0.7 | 0.57 | 0.013 | 0.034 | 0.42 | 0.048 |
| LTB4 | 0.3 | 0.41 | 0.6 | 0.64 | 0.67 | 0.67 | 0.014 |
| 8iso_PGA1 | 0.3 | 0.33 | 0.23 | 0.22 | 0.82 | 0.36 | 0.32 |
| 5_iPF2a_VI | 0.3 | **0.012** | 0.31 | 0.01 | 0.94 | 0.35 | 0.59 |
| 9_HOTrE | 0.31 | 0.075 | 0.61 | 0.008 | 0.79 | 0.014 | 0.23 |
| (TCDCA+GCDCA)/CDCA | 0.31 | 0.82 | 0.44 | 0.22 | 0.27 | 0.011 | 0.92 |
| 9-HODE/LA | 0.32 | 0.35 | 0.87 | 0.07 | 0.23 | 0.3 | 0.66 |
| LPE20.3 | 0.32 | 0.33 | 0.46 | 0.89 | 0.67 | 0.012 | 0.92 |
| TCDCA/CDCA | 0.32 | 0.82 | 0.44 | 0.22 | 0.27 | 0.01 | 0.91 |

| | | | | | | | |
|---|---|---|---|---|---|---|---|
| ResolvinE2/EPA | 0.32 | 0.77 | 0.92 | 0.14 | 0.011 | 0.027 | 0.21 |
| LPE22.5 | 0.32 | 0.47 | 0.95 | 0.015 | 0.84 | 0.0053 | 0.089 |
| FA18.2_w6 | 0.33 | **0.0088** | 0.87 | 0.2 | 0.59 | <.0001 | 0.098 |
| thromboxane_B2 | 0.33 | 0.73 | 0.99 | 0.55 | 0.5 | 0.67 | 0.067 |
| LPE18.0 | 0.34 | 0.67 | 0.94 | <.0001 | 0.096 | 0.0017 | 0.46 |
| LPE16.0 | 0.35 | 0.85 | 0.85 | <.0001 | 0.87 | 0.0003 | 0.048 |
| FA18.3_w3 | 0.36 | 0.41 | 0.86 | 0.25 | 0.049 | 0.025 | 0.41 |
| 20-HETE/AA | 0.37 | 0.95 | 0.82 | 0.054 | 0.94 | 0.03 | 0.27 |
| PGA2/AA | 0.38 | 0.41 | 0.54 | 0.28 | 0.49 | 0.0006 | 0.6 |
| 12-HEPE/12-HETE | 0.38 | 0.85 | 0.29 | 0.0032 | 0.2 | 0.029 | 0.0024 |
| LPI18.2 | 0.38 | 0.29 | 0.82 | 0.89 | 0.13 | 0.48 | <.0001 |
| 12_13_EpOME | 0.38 | 0.28 | 0.67 | 0.0021 | 0.051 | 0.087 | 0.84 |
| FA18.3_w6 | 0.39 | **0.013** | 1 | 0.58 | 0.16 | <.0001 | 0.25 |
| PGE3/EPA | 0.39 | 0.28 | 0.34 | 0.87 | 0.29 | 0.0033 | 0.21 |
| FA22.5_w6 | 0.4 | 0.12 | 0.47 | 0.57 | 0.78 | <.0001 | 0.52 |
| GLCA/LCA | 0.4 | 0.21 | 0.7 | 0.32 | 0.15 | 0.65 | 0.79 |
| LPA14.0 | 0.4 | 0.41 | 0.87 | 0.034 | 0.82 | <.0001 | 0.26 |
| 1_OG_2_OG | 0.41 | 0.29 | 0.65 | 0.68 | 0.15 | 0.23 | 0.37 |
| LPG22.4 | 0.42 | 0.47 | 0.14 | 0.036 | 0.01 | 0.83 | 0.046 |
| 12_13_DiHOME | 0.42 | 0.94 | 0.48 | 0.0001 | 0.015 | 0.0005 | 0.47 |
| LPG18.0 | 0.43 | 0.053 | 0.3 | 0.007 | 0.77 | 0.8 | 0.018 |
| LPI18.0 | 0.43 | 0.05 | 0.69 | 0.11 | 0.24 | <.0001 | 0.0069 |
| LPI20.4 | 0.43 | **0.043** | 0.23 | 0.19 | 0.048 | 0.084 | 0.5 |
| LPG22.6 | 0.43 | 0.67 | 0.98 | 0.017 | 0.0011 | 0.31 | 0.098 |
| 5-HEPE/5-HETE | 0.44 | 0.65 | 0.24 | 0.082 | 0.016 | 0.0015 | 0.018 |
| 5_HETE | 0.44 | 0.69 | 0.32 | 0.45 | 0.34 | <.0001 | 0.29 |
| 8iso_PGA2 | 0.45 | 0.98 | 0.13 | 0.006 | 0.027 | 0.0014 | 0.017 |
| ResolvinE2 | 0.45 | 0.79 | 0.92 | 0.98 | 0.24 | 0.56 | 0.85 |
| 5_iPF2a_VI/AA | 0.46 | **0.0067** | 0.18 | 0.066 | 0.87 | 0.13 | 0.52 |
| 9_KODE | 0.46 | 0.067 | 0.85 | 0.28 | 0.28 | 0.0042 | 0.21 |
| LPE18.2 | 0.47 | 0.076 | 0.91 | 0.003 | 0.14 | 0.045 | 0.16 |
| LPE14.0 | 0.48 | 0.16 | 0.98 | 0.52 | 0.1 | 0.023 | 0.088 |
| TCA | 0.48 | 0.29 | 0.07 | 0.17 | 0.78 | 0.58 | 0.78 |
| TDCA/DCA | 0.49 | 0.33 | 0.86 | 0.16 | 0.58 | 0.057 | 0.97 |
| 11-HETE/AA | 0.5 | 0.59 | 0.39 | 0.7 | 0.48 | 0.74 | 0.47 |
| 12_HHTrE | 0.51 | 0.71 | 0.98 | 0.76 | 0.39 | 0.75 | 0.1 |
| 5-HETE/AA | 0.51 | 0.8 | 0.86 | 0.67 | 0.67 | 0.87 | 0.24 |
| (TDCA+GDCA)/DCA | 0.51 | 0.38 | 0.91 | 0.14 | 0.93 | 0.064 | 0.89 |
| LPS22.6 | 0.52 | 0.053 | 0.58 | 0.39 | 0.15 | 0.49 | 0.79 |
| 15_HETE | 0.52 | 0.86 | 0.56 | 0.95 | 0.062 | 0.13 | 0.22 |
| 12_13_EpOME/LA | 0.52 | 0.22 | 0.65 | 0.011 | 0.023 | 0.17 | 0.76 |
| GUDCA | 0.53 | 0.5 | 0.15 | 0.92 | 0.097 | 0.93 | 0.31 |
| Sphingosine1P18.2 | 0.53 | 0.36 | 0.35 | <.0001 | 0.24 | <.0001 | 0.021 |
| LPA18.2 | 0.54 | **0.003** | 0.95 | 0.0062 | 0.83 | <.0001 | 0.22 |
| FA20.3_w9 | 0.54 | 0.44 | 0.81 | 0.97 | 0.44 | <.0001 | 0.39 |

| | | | | | | | |
|---|---|---|---|---|---|---|---|
| LPG20.4 | 0.55 | 0.97 | 0.15 | 0.2 | 0.082 | 0.29 | 0.043 |
| 8,9-DiHETrE/AA | 0.57 | 0.62 | 0.86 | 0.18 | 0.95 | 0.014 | 0.33 |
| LPG20.3 | 0.58 | 0.58 | 0.13 | 0.027 | 0.25 | 0.32 | 0.4 |
| 15-HETE/AA | 0.59 | 0.56 | 0.75 | 0.83 | 0.12 | 0.17 | 0.11 |
| 12_13_DiHOME/LA | 0.61 | 0.3 | 0.6 | 0.0021 | 0.0044 | 0.2 | 0.87 |
| 13-HODE/LA | 0.62 | 0.3 | 0.54 | 0.013 | 0.13 | 0.69 | 0.59 |
| LPE20.5 | 0.62 | 0.14 | 0.6 | 0.0004 | 0.04 | 0.0018 | 0.11 |
| delta17_6_keto_PGF1a/EPA | 0.62 | 0.56 | 0.56 | 0.22 | 0.33 | 0.054 | 0.9 |
| GCDCA/GLCA | 0.63 | 0.38 | 0.66 | 0.85 | 0.99 | 0.088 | 0.61 |
| 5-HEPE/EPA | 0.64 | 0.55 | 0.79 | 0.0003 | 0.015 | 0.31 | 0.73 |
| cLPA16.1 | 0.64 | 0.95 | 0.26 | 0.096 | 0.8 | 0.0003 | 0.12 |
| LPG18.1 | 0.65 | 0.11 | 0.34 | 0.011 | 0.81 | 0.033 | 0.89 |
| LPG18.2 | 0.65 | 0.28 | 0.29 | 0.46 | 0.18 | 0.31 | 0.29 |
| GUDCA/UDCA | 0.66 | 0.55 | 0.69 | 0.55 | 0.22 | 0.35 | 0.95 |
| AEA | 0.67 | 0.24 | 0.19 | 0.63 | 0.25 | 0.088 | 0.73 |
| LTE4 | 0.67 | 0.88 | 0.3 | 0.72 | 0.28 | 0.029 | 0.73 |
| 9_HODE | 0.68 | 0.71 | 0.76 | 0.019 | 0.46 | 0.0002 | 0.88 |
| (GLCA+TLCA)/LCA | 0.68 | 0.62 | 0.94 | 0.4 | 0.42 | 0.47 | 0.81 |
| LPE22.4 | 0.69 | 0.72 | 0.41 | 0.61 | 0.088 | 0.17 | 0.038 |
| 8iso_PGF3a | 0.69 | 0.84 | 0.51 | 0.4 | 0.04 | 0.52 | 0.029 |
| thromboxane_B3 | 0.7 | 0.88 | 0.84 | 0.093 | 0.045 | 0.93 | 0.93 |
| LPG14.0 | 0.71 | **0.022** | 0.7 | 0.0037 | 0.49 | 0.24 | 0.017 |
| 11_HETE | 0.72 | 0.99 | 0.86 | 0.59 | 0.39 | 0.0051 | 0.52 |
| LPG16.0 | 0.73 | 0.054 | 0.66 | 0.015 | 0.15 | 0.5 | 0.12 |
| LPE16.1 | 0.73 | 0.25 | 0.81 | 0.017 | 0.82 | <.0001 | 0.94 |
| (TCA+GCA+TDCA+GDCA)/(GUDCA+GLCA+TLCA+TCDCA+GCDCA) | 0.73 | 0.43 | 0.24 | 0.39 | 0.61 | 0.34 | 0.47 |
| 12_13_DiHOME/12_13_EpOME | 0.74 | 0.15 | 0.82 | 0.81 | 0.7 | 0.2 | 0.65 |
| LPI22.4 | 0.75 | 0.24 | 0.38 | 0.0075 | 0.23 | 0.5 | 0.29 |
| TLCA/LCA | 0.77 | 0.73 | 0.61 | 0.32 | 0.71 | 0.088 | 1 |
| delta17_6_keto_PGF1a | 0.79 | 0.52 | 0.45 | 0.88 | 0.82 | 0.91 | 0.64 |
| PGE3 | 0.82 | 0.24 | 0.34 | 0.016 | 0.32 | 0.9 | 0.98 |
| DCA/(LCA+UDCA) | 0.82 | 0.34 | 0.79 | 0.073 | 0.46 | 0.77 | 0.16 |
| sum_HODE | 0.84 | 0.62 | 0.62 | 0.0075 | 0.36 | <.0001 | 0.55 |
| LPE18.3 | 0.86 | 0.19 | 0.36 | 0.0001 | 0.0048 | 0.68 | 0.91 |
| LPE18.1 | 0.87 | 0.094 | 0.49 | 0.016 | 0.75 | 0.74 | 0.19 |
| (GDCA+GLCA)/(TDCA+TLCA) | 0.87 | 0.1 | 0.28 | 0.81 | 0.23 | 0.011 | 0.77 |
| CDCA | 0.88 | 0.53 | 0.038 | 0.053 | 0.62 | 0.04 | 0.76 |
| 13_HODE | 0.89 | 0.5 | 0.46 | 0.0022 | 0.32 | <.0001 | 0.26 |
| 5_HETrE | 0.91 | 0.78 | 0.42 | 0.76 | 0.33 | 0.0022 | 0.47 |
| sum_HETrE | 0.92 | 0.9 | 0.72 | 0.82 | 0.71 | 0.078 | 0.16 |
| HCA | 0.94 | 0.072 | 0.17 | 0.67 | 0.5 | 0.079 | 0.45 |
| 17-HDoHE/DHA | 0.94 | 0.13 | 0.39 | 0.75 | 0.51 | 0.008 | 0.86 |
| LPG18.3 | 0.94 | 0.14 | 0.038 | 0.39 | 0.13 | 0.91 | 0.65 |

| | | | | | | | |
|---|---|---|---|---|---|---|---|
| Sphingosine1P16.1 | 0.94 | 0.46 | 0.65 | <.0001 | 0.75 | 0.001 | 0.0061 |
| LPI18.1 | 0.95 | **0.0056** | 0.21 | 0.64 | 0.83 | 0.069 | 0.57 |
| GDCA/GLCA | 0.95 | 0.72 | 0.22 | 0.2 | 0.27 | 0.86 | 0.51 |
| 9-HEPE/EPA | 0.96 | 0.31 | 0.76 | 0.062 | 0.0003 | 0.45 | 0.66 |
| 5,6-DiHETrE/AA | 0.96 | 0.59 | 0.62 | 0.044 | 0.31 | 0.067 | 0.065 |
| TDCA/TLCA | 0.96 | 0.91 | 0.93 | 0.29 | 0.84 | 0.41 | 0.24 |
| 15_HETrE | 0.97 | 0.88 | 0.76 | 0.66 | 0.38 | 0.15 | 0.45 |
| cLPA14.0 | 0.98 | 0.23 | 0.79 | 0.056 | 0.57 | 0.38 | 0.0003 |
